# Does offspring demand affect parental life-history trade-offs?

**DOI:** 10.1101/2020.09.09.289298

**Authors:** M.I. Mäenpää, P.T. Smiseth

## Abstract

Life-history trade-offs between the number and size of offspring produced, and the costs of reproduction on future reproduction and survival can all be affected by different levels of parental effort. Because of these trade-offs the parents and the offspring have different optima for the amount of care given to the current brood, which leads to a conflict between parents and offspring. The offspring, as well as the parents, have the ability to affect parental effort, and thus changes in offspring traits have the potential to cause reproductive costs on the parents. Here, we used a repeated cross-fostering design to manipulate offspring demand during juvenile development in the burying beetle *Nicrophorus vespilloides* to examine whether responding to offspring begging incurs reproductive costs to the parent. After a manipulated first reproductive event, we gave each experimental female, that had been exposed to different levels of offspring demand, a chance to breed again, and monitored their survival. We found that larval demand influences the trade-off between the size and number of offspring produced, but has no impact on the reproductive costs through future reproduction or survival of the parent. The parents do, however, pay an overall fecundity cost for the general success of their first broods, but this cost was not related to the changes in the levels of larval begging. Other traits, including survival showed no costs of reproduction. Survival and the number of larvae successfully raised in the second broods correlated positively, indicating differences in the individual quality of the parents.

## Introduction

Maximising parental fitness during multiple reproductive events requires successful optimisation of resource allocation between current and future reproduction, as well as survival (Stearns, 1992). Within broods, the parents face a trade-off between the number and size of offspring produced (Lloyd, 1987; Smith and Fretwell, 1974), and between broods, they must balance the reproductive costs (as first proposed by Williams, 1966) that caring for the current brood incurs for their future reproduction (eg. Gustafsson and Sutherland, 1988; Monaghan and Nager, 1997), or survival (eg. Dijkstra et al., 1990). Previous work investigating costs of reproduction have included observational studies on correlations between life-history traits (eg. Bryant, 1979), and experimental studies that have manipulated parental effort through traits related to current reproduction (such as clutch size, eg. Hodges et al., 2015; Kölliker et al., 2015), or by adding to the energetic costs of care for the parents (such as handicapping the parent with weights (eg. Harding et al., 2009; Hegemann et al., 2013; Tieleman et al., 2008). The evidence for these life-history trade-offs still remains controversial, as many empirical studies have not found the expected trade-off between the life-history traits, or have even found a positive relationship instead of the expected negative correlation (reviewed in Reznick, 1985; Roff and Fairbairn, 2007). The lack of phenotypic trade-off in some studies can be due to quality differences between the individuals used in the study (ie. among-individual heterogeneity; Weladji et al., 2008, but see Wilson and Nussey, 2010 for critique regarding the terminology), or for example unaccounted for sex differences (see Santos and Nakagawa, 2012). In addition, trade-offs discovered through inducing different levels of parental effort have so far ignored the potential for parental effort to change based on the traits and the behaviours of the offspring themselves, the potential for which has been reported widely in other contexts (Agrawal et al., 2001; Kilner and Johnstone, 1997; Parker et al., 2002; Rodríguez-Gironés et al., 2001; Smiseth et al., 2003). Aside from manipulations of offspring number, no studies have been conducted to test the effects of manipulating the offspring’s behaviour rather than the costs and benefits of care to the parent.

Offspring influence over parental care relies on behavioural plasticity of the parent in responding to the cues received from the offspring, as offspring demand has an effect on parental supply (Hussell, 1988). Offspring induced plasticity has been reported before: for example, Kight (1997) found that the duration of care and parental defence was affected by cues from the offspring, which were eventually overridden by the parent’s internal clock in the burrower bug *Sehirus cinctus*. Theoretically this plasticity is only beneficial for the parent when the benefits from a unit of care given are higher than benefits from investing the same unit into future reproduction. Furthermore, the gains from future reproduction change based on parent’s likelihood of breeding again. Indeed, Thorogood et al. (2011) found that the parents changed their responsiveness to offspring food solicitation signals when their potential to breed again was manipulated through food supplementation, in a New Zealand passerine, *Notiomystis cincta*. Therefore, as the parents are trying to optimise their care across multiple broods, and the offspring benefit from each unit of investment they receive, the optimal levels of care are different for the parents and offspring, which leads to a conflict between the parents and the offspring (Trivers, 1974). Parent-offspring conflict has the potential to sway the trade-offs between parental effort and life-history traits to a direction determined not only by the resource allocation constraints affecting the parent, but also by the need of the offspring. Both the parents and the offspring can bias the amount of parental investment towards their own optimum through influencing one another by communicating (reviewed in Kilner and Johnstone, 1997). This communication has in many taxa evolved into elaborate begging displays used by the offspring to elicit food from the parents, and to give the parent honest information about the condition of the offspring (Kilner and Johnstone, 1997). To optimise the amount of care given to the offspring, the parents not only need to assess their potential to breed again, but also assess the potential pay-off they can receive from their current offspring based on their communication cues.

In this study, we experimentally manipulated offspring demand during the first reproductive event to investigate whether offspring demand can incur a reproductive cost to the parent. The burying beetle, *Nicrophorus vespilloides*, is an excellent system for such experiments, as the larvae of the species exhibit begging behaviour, the intensity of which is determined by larval age (Smiseth et al., 2003), and as such can be manipulated with ease (Mäenpää & Smiseth, in press). Burying beetles feed and breed on carrion of small vertebrates and provide post-hatching parental care for the offspring (Eggert et al., 1998; Scott, 1998). Usually only the females stay with the brood until the larvae disperse from the carcass into the soil to pupate (Scott, 1998). The larvae are capable of feeding from a suitably prepared carcass on their own (Eggert et al., 1998), but their growth and survival are positively affected by parental food provisioning (Smiseth and Moore, 2004). Larvae beg by touching the adults with their feet (Smiseth et al., 2003; Smiseth and Moore, 2002), and the parents respond by provisioning pre-digested carrion to their broods (Scott, 1998). As the larvae become more proficient in self-feeding during juvenile period, the intensity of begging in a brood declines, leading to age-dependent variation in the amount of begging exhibited by an individual larva (Smiseth et al., 2003). Begging peaks at 24 hours after hatching, and starts declining after that, until approximately 72 hours after hatching, which marks the point of transitioning to nutritional independence (Smiseth et al., 2003). Here, we manipulated the age of the brood female burying beetles were caring for during their first reproductive event, in order to manipulate the level of offspring demand the females experienced. Our aim was to test whether the changes in larval demand affected the trade-off between size and number of offspring produced, and whether the female parents would exhibit cumulative reproductive costs associated with responding to the manipulated demand. We predict that changes in larval demand would lead to changes at different stages in the life-history of the parent. Specifically, (1) the number of offspring produced in the second brood would be expected to increase or decrease depending on the levels of begging experienced in the first broods, similarly to (2) the success of the second brood, and (3) the survival of the parent.

## Materials and Methods

### Origin and husbandry of the beetles

The beetles used in this experiment derived from a large, outbred laboratory population originating from wild-caught beetles trapped in Corstophine Hill and Craiglockhart Hill (Edinburgh, UK), Kennall Vale, (Cornwall, UK) and Madingley Woods (Cambridge, UK). All beetles in the laboratory population were kept individually in transparent plastic containers (7 x 12 x 6 cm) under constant light at 20°C, and fed small pieces of organic beef twice a week.

### Experimental treatments: first broods

We randomly selected pairs of nonsibling virgin male and female beetles to be mated. All matings were conducted in transparent containers (12 x 18 x 6 cm) filled with 2 cm of moist soil and a previously frozen mouse carcass to breed on (range 20-25 g, supplied by Livefoods Direct Ltd, Sheffield, UK). We only used female parents in this experiment to remove the confounding effects of sex-differences in the expression of the life-history trade-offs, and chose females because male care is highly variable and has no detectable effects on larval growth or survival under laboratory conditions (Eggert et al., 1998; Smiseth et al., 2005). Thus, we removed the male 60 hours after pairing, which is before the larvae started hatching. Concurrently, in order to have control over the age of the experimental broods and prevent any contact between the female and the larvae prior to brood age manipulations, we moved the female and the carcass into a new container filled with soil to separate the eggs from the breeding female. The egg boxes were checked 5 times a day for hatching. When the larvae started hatching, we assigned some females into treatments, and used others to create donor broods for the brood age manipulations. The donor females were assigned mixed maternity broods of 15-25 newly hatched larvae from the supply of broods that had started hatching in the egg boxes. These donor broods were then used to create experimental broods after they had reached an age appropriate for the treatment (described below) of the experimental female they were given to.

We manipulated larval demand in the experimental broods by repeated cross-fostering for the approximate duration of larval dependency (i.e. first 72 hours after hatching). We gave each experimental female a brood of 10 mixed maternity larvae of a known age, and swapped the brood with another experimental brood of a known age systematically throughout the experiment, using a supply of donor broods consisting of larvae of an appropriate age. We generated four treatments that differed with respect to the ages of the broods. (1) The broods of the control females were initially set up with newly hatched larvae, and the broods were always swapped with a brood consisting of larvae of the same age as the ones taken away. This control group was created to control for the effects of handling on larval and parental behaviours. (2) To create a treatment with high larval demand (hereafter referred to as the high demand treatment), we kept the larvae young throughout the manipulation period, by supplying the female with a brood of 1-hour-old larvae, which were always swapped as they reached the age of 25 hours to another brood of 1-hour-old larvae. (3) In the intermediate demand treatment, the female was given 25-hour-old larvae, which were swapped at the age of 49 hours with another brood of 25-hour-old larvae. (4) Finally, in the low demand treatment, the female was given 49-hour-old larvae, which were swapped at the age of 73 hours with another brood of 49-hour-old larvae. In all treatments, the larvae were swapped four times in total during the manipulation period. To capture the initial behaviour of the females when given larvae of a different age than expected (data presented in Mäenpää & Smiseth, in press), the first swaps were conducted an hour after the initial broods were given to the females. To ensure that the age of these first broods corresponded to the rest of the experimental manipulations, the first broods given to the females consisted of newly hatched, 24-hour-old, 48-hour-old, and 72-hour-old larvae in the control, high, intermediate, and low demand treatments, respectively. An hour later these broods were swapped, and the swaps were then continued as determined previously. After the last swap, the female was allowed to raise the larvae until they dispersed from the carcass.

When the brood had dispersed, i.e. when all larvae had moved from the carcass to the soil around it, the female was returned to individual containment. We then weighed the broods to the nearest 0.1 mg using a digital scale (Ohaus Pioneer, with 0.1 mg accuracy), and counted the number of surviving larvae. Due to mortality in the donor broods, experimental females were occasionally discarded in the middle of the experiment, as there were no larvae to provide for them. Thus, our final sample sizes for the control treatment, and the high, intermediate, and low demand treatments were 20, 27, 19 and 18, respectively.

### Recording reproductive costs: second broods and lifespan

To record reproductive costs of caring for broods with different levels of larval demand, we mated the experimental females for a second time to another unrelated virgin male, 3-11 days after the dispersal of the first broods. The males were again removed 60 hours after hatching to remove the confounding effects of male care. We then monitored the unmanipulated second broods to determine the reproductive success of the females. To assess fecundity, we counted the eggs laid at the bottom of the containers, visible through the transparent plastic, which correlates strongly with the total number of eggs laid (Monteith et al., 2012). We checked the boxes daily first for hatching, and then for dispersal, as well as the death of the female or the brood. At dispersal, we weighed the broods and counted the number of larvae, to assess reproductive success in the number and size of larvae that the females successfully raised. After the larvae had dispersed, we moved the females to individual housing in smaller containers. The boxes were checked daily for the death of the females. The recorded day of death was then used to determine the lifespan of an experimental female in days, starting from the day it had eclosed as an adult. Four females in the high demand treatment died before they could be mated again, and three beetles escaped their containers during mortality tracking, leaving our final sample size on survival data with 19, 26, 19 and 17 for the control, high, intermediate and low treatments, respectively.

### Statistical analyses

All analyses were conducted with R version 3.4.1 (R Core Team, 2017). We used linear mixed effects models (package lme4, Bates et al., 2014) as all response factors (individual larva mass at dispersal, number of eggs, number of larvae at dispersal, and lifespan) had a gaussian error distribution. In all analyses, we included experimental block as a random variable.

First, we explored the trade-off between number and size of offspring within different treatments in both first and second broods. We assigned average (individual) larva mass at dispersal as the response variable, and the brood size at dispersal as the explanatory variable. Treatment (control, high, intermediate and low), brood number (first, second) were then added as fixed factors. We included three two-way interaction terms in the model: (1) The interaction between treatment and brood size was added to determine whether the potential trade-off curve differed in shape between treatments. (2) The interaction between brood number and brood size was included to explore whether the shape of the trade-off curve differed between first and second brood. (3) The interaction between treatment and brood number was added to determine whether the average size of larvae within a treatment changed differently going from first to the second broods. To explore the overall shape of the trade-off curve, we also conducted a separate mixed effects model, where we used the average (individual) larva mass at dispersal as the response variable, and the brood size at dispersal and its quadratic term as the explanatory variables. To avoid pseudorepliction, we included female identity as an additional random effect in both of these models.

Next, we investigated the reproductive costs that changes in offspring demand could have imposed upon the female parent using separate models for the traits we predicted them to be found in: (1) number of eggs laid, (2) number of larvae at dispersal, and (3) lifespan of the female. In all these models, the Treatment (control, high, intermediate, low) and the size of the first brood at disperal were added to the models as fixed factors, to explore the effects of offspring demand, and overall reproductive success at the first reproductive event, respectively. We also included the two-way interaction between the two variables to explore whether the effect of treatment could impose a reproductive cost through the overall reproductive success of the first brood. In the lifespan model, we also included the size of the female’s second brood (at dispersal), and its two-way interaction with Treatment, as an explanatory variable, as the trade-off between reproduction and survival may also act via the success of the second brood, which may have different effects in different treatments.

To account for effects of female age, which can be a strong determinant for traits associated with the success of offspring produced (Kindsvater and Alonzo, 2014), we age at first reproduction as a covariate in all models. Our data had a disproportionate number of young females in the different treatments. Thus, when female age had a significant effect, we repeated the analyses with a subset of the data only using the females that were older than 20 days at the time of the first reproduction to test for the robustness of the results. In this occasion, our sample sizes were 17, 16, 13 and 16, for the control, high, intermediate and low treatments, respectively. As we found no differences from the overall main results, we only report the results of the original models with full data. We also assigned mouse masses of the first and second reproductive events as covariates in all models, as the amount of resources available affects the number and size of offspring produced (Smiseth and Moore, 2002). To attain parsimonious models, non-significant (P<0.1) terms were removed based on ANOVA’s comparing the maximum likelihood estimates of the nested models.

## Results

### Number and size of offspring

The overall relationship between number and size of offspring was quadratic in shape (Figure 1a), with a negative quadratic term (coef(SE)= −1.15 x 10^−4^(0.22 x 10^−4^), F_1, 142_= 27.08, *P* <0.001), and a positive linear term (coef(SE)= 3.85 x 10^−3^(0.91 x 10^−3^), F_1, 143_= 17.92, *P* <0.001). Larval mass increased up to brood sizes of approximately 15 larvae, and decreased after that (Figure 1a), indicating that the trade-off between the number and size of the offspring was only apparent in larger brood sizes.

**Figure 1:**
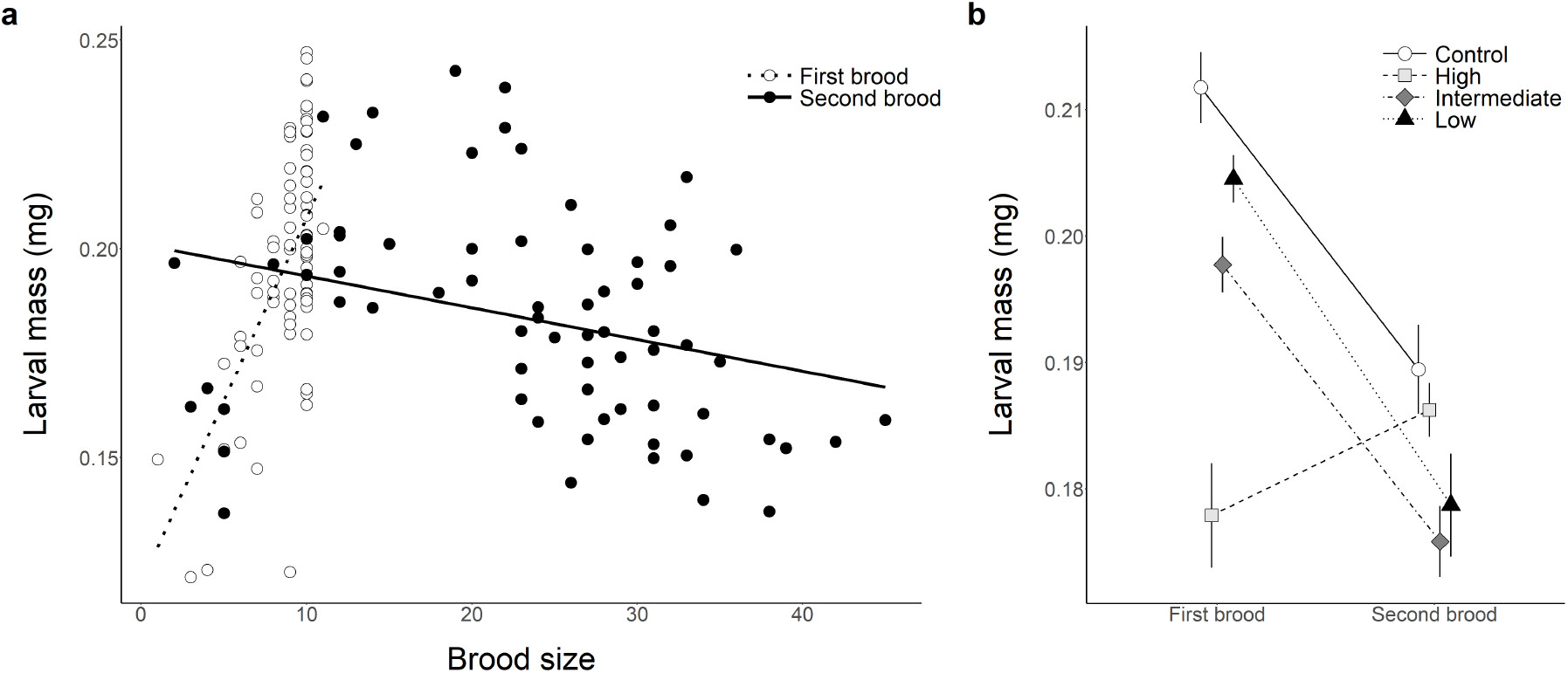
Size and number of offspring in manipulated first broods, and the unmanipulated second broods of the female burying beetles (*Nicrophorus vespilloides*). The number of larvae was kept at a maximum of 10 larvae that were not related to the females in the first broods, while it was allowed to vary freely in the biological second broods. The treatments indicate different levels of manipulated larval demand. (a) Individual body mass (brood means) at dispersal in relation to brood size. (b) Individual body mass (mean ± SE) at dispersal for the larvae produced in the first and second reproductive events.

Larval demand, manipulated through different treatments, affected the relationship between the number and size of the offspring produced (Treatment: Brood size interaction in Table 1). This effect was due to the high demand treatment differing from the other treatments (Table 1), and showing overall a positive relationship between the variables, while other treatments exhibited the expected trade-off curve. The females in the high demand treatment produced larger offspring in their second brood unlike the females of the other treatments, which produces smaller offspring in their second broods (Figure 1b). Overall, there was a reversal in the linear relationship between number and size of the offsrpring in the manipulated first broods, which showed a positive association, and the unmanipulated second broods, for which the relationship was negative (Brood number: Brood size interaction in Table 1, Figure 1a). This result was likely due to the experimental manipulations limiting the number of larvae in the first broods to a maximum of 10 larvae, which could have indicated that the broods were not large enough for the trade-off to take place. Generally, the size of the larvae in the first broods increased with decreasing larval demand (see treatment order in Figure 1b, Table 1). The unmanipulated second broods consisted of more larvae (Figure 1a), that were on average more closely matched in size in the different treatments than the larvae of the first broods (Figure 1b). The size of the carcass also had a positive effect on the size of the offspring (Table 1).

**Table 1:** Factors affecting the size of offspring in two reproductive events of the burying beetle *Nicrophorus vespilloides*. All estimates are derived from a linear mixed effects model (lmer) with experimental block and the identity of the female assigned as random factors. Degrees of freedom and *P*-values attained through Satterwaithe approximation. We present parameter estimates, standard errors, t-statistics, and *P*-values for each factor level, as well as *χ*^2^ statistics and *P*-values for the overall effects for factors (in bold).

| Variable | Par | SE | df | $\chi^2/t$ | *P* |
| --- | --- | --- | --- | --- | --- |
| Treatment |  |  | **3, 128** | **3.58** | **0.016** |
| High | -0.03 | 0.01 | 132 | -3.09 | 0.002 |
| Intermediate | -0.01 | 0.01 | 128 | -1.00 | 0.318 |
| Low | -0.01 | 0.01 | 127 | -1.26 | 0.215 |
| Brood number |  |  | **1, 131** | **13.12** | **<0.001** |
| Second brood | 0.06 | 0.02 | 131 | 3.62 | <0.001 |
| Brood size | 0.01 | 0.002 | 132 | 2.62 | 0.010 |
| Mouse mass | 0.003 | 0.001 | 133 | 2.33 | 0.021 |
| Treatment: Brood number* |  |  | **3, 93** | **1.00** | **0.399** |
| High: Seond brood | -0.003 | 0.014 | 93 | -0.22 | 0.828 |
| Intermediate: Second brood | 0.029 | 0.020 | 92 | 1.46 | 0.147 |
| Low: Second brood | 0.012 | 0.016 | 84 | 0.74 | 0.459 |
| Treatment: Brood size |  |  | **3, 101** | **3.92** | **0.011** |
| High: Brood size | 0.002 | 0.001 | 116 | 2.94 | 0.004 |
| Intermediate: Brood size | 0.0004 | 0.001 | 106 | 0.76 | 0.449 |
| Low: Brood size | 0.001 | 0.001 | 102 | 0.91 | 0.365 |
| Brood number: Brood size |  |  | **1, 132** | **15.35** | **<0.001** |
| Second brood: Brood size | -0.007 | 0.002 | 132 | -3.92 | <0.001 |

\* Factor removed in the model simplification process. Estimates derived from the last model the factor was included in.

### Reproductive costs

Manipulated larval demand (Treatment) did not affect any of the traits likely to show the costs of reproduction (Figure 2a-c, Table 2). However, the number of eggs produced in the second broods declined significantly with each surviving offspring of the first broods (Figure 2d, Table 2). This effect remained even after repeating the analysis after removing the single brood that contained no surviving larvae in the first reproductive event. Thus, the parents paid a fecundity cost for the general success of their first broods. No such effects were detected for the nuber of offspring surviving to dispersal in the second broods (Figure 2e, Table 2). However, the number of eggs produced was a strong predictor for the number of offspring subsequently surviving to dispersal from the broods (lm, *β*(SE)= 0.65(0.08), R^2^= 0.48, F_1,67_= 62.41, *P* <0.001), indicating an indirect effect of success of the first brood on the success of the second brood as well. The lifespan of the females also showed no indication of reproductive costs (Figure 2f, Table 2). There was, however, a positive effect of the number of larvae dispersing from the second broods (Table 2), indicating that the females producing larger second broods also had a longer lifespan.

**Figure 2:**
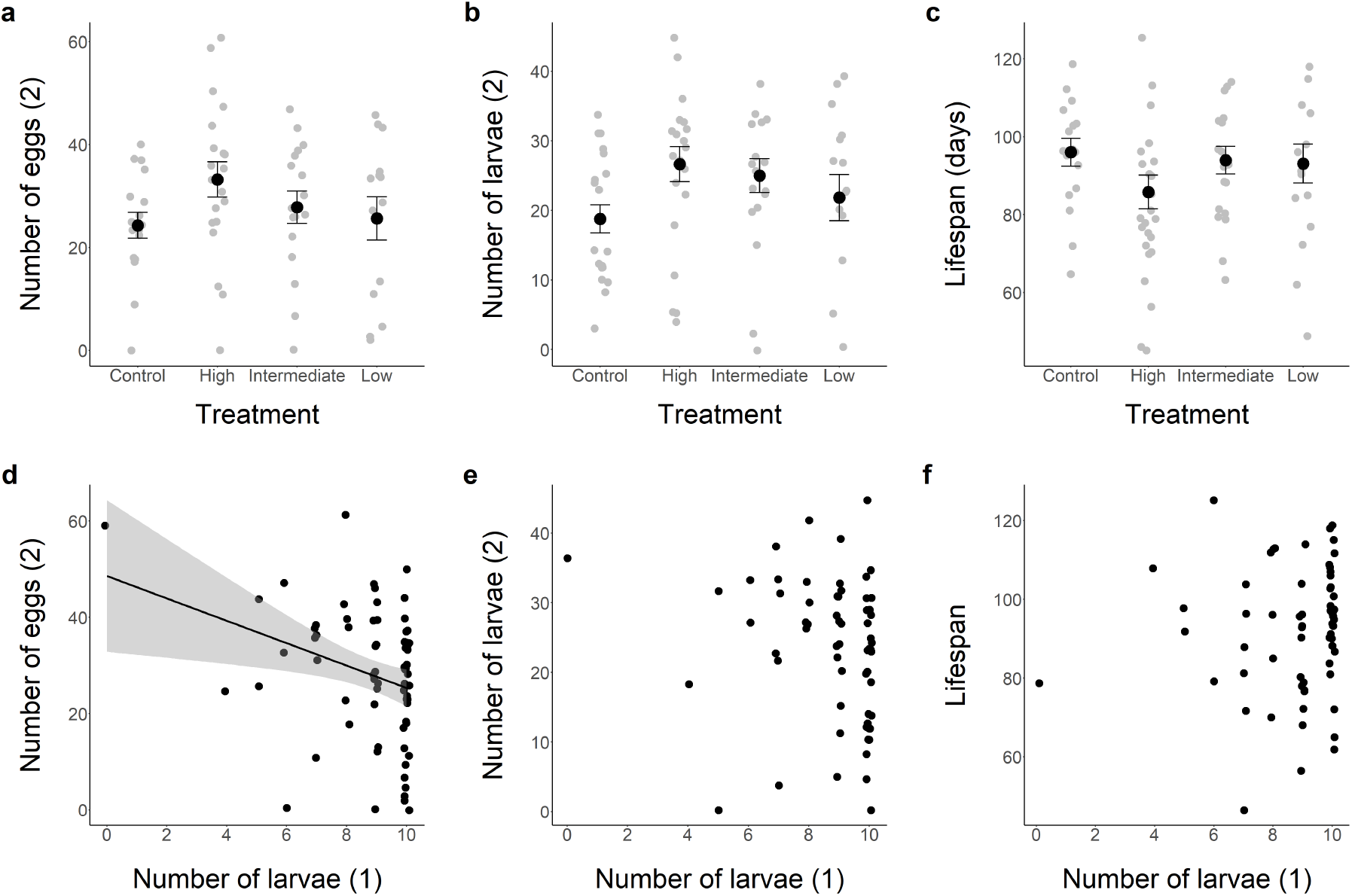
The effects of treatments manipulating larval demand (a-c), and the general success of the first broods (d-f) on traits likely to show costs of reproduction (number of eggs (a, d), number of larvae at dispersal (b, e), and lifespan (c, f)) in *Nicrophorus vespilloides*. The number indicated in brackets identifies the reproductive event for which the trait was measured (1= first broods, 2= second broods). The grey areas indicate standard errors for the regression lines.

**Table 2:** Factors affecting the life-history traits likely to show reproductive costs in female burying beetles (*Nicrophorus vespilloides*). All estimates are derived from a linear mixed effects models (lmer), with degrees of freedom and P-values attained through Satterwaithe approximation. The number of the brood is indicated in brackets after each variable that was measured in both reproductive events (1= first broods, 2= second broods).

| Response | Variable | Par | SE | df | $\chi^2/t$ | P |
| --- | --- | --- | --- | --- | --- | --- |
| Number of eggs (2) | Treatment* |  |  | **3, 61** | **0.02** | **0.996** |
|  | High | 0.49 | 5.41 | 61 | 0.09 | 0.928 |
|  | Intermediate | 0.85 | 4.82 | 61 | 0.18 | 0.861 |
|  | Low | -0.40 | 4.73 | 61 | -0.09 | 0.932 |
|  | Brood size (1) | -1.95 | 0.88 | 65 | -2.21 | 0.030 |
|  | Female age | -0.16 | 0.10 | 65 | -1.65 | 0.104 |
|  | Mouse mass | 2.00 | 1.15 | 65 | 1.74 | 0.087 |
| Brood size<br>at dispersal (2) | Treatment |  |  | **3, 54** | **2.01** | **0.123** |
|  | High | -35.08 | 93.82 | 54 | -0.37 | 0.710 |
|  | Intermediate | -61.66 | 94.84 | 54 | -0.65 | 0.518 |
|  | Low | 71.53 | 110.85 | 54 | -0.65 | 0.521 |
|  | Brood size (1) | -4.06 | 9.42 | 54 | -0.43 | 0.668 |
|  | Female age | -0.24 | 0.08 | 54 | -3.06 | 0.003 |
|  | Treatment: Brood size (1) |  |  | **3, 54** | **2.11** | **0.110** |
|  | High | 3.85 | 9.46 | 54 | 0.41 | 0.686 |
|  | Intermediate | 7.02 | 9.58 | 54 | 0.73 | 0.467 |
|  | Low | -7.29 | 11.21 | 54 | -0.65 | 0.518 |
| Lifespan | Treatment* |  |  | **3, 52** | **1.05** | **0.380** |
|  | High | -7.57 | 5.61 | 55 | -1.35 | 0.183 |
|  | Intermediate | -0.47 | 5.11 | 55 | -0.09 | 0.926 |
|  | Low | 0.36 | 5.02 | 55 | 0.07 | 0.943 |
|  | Brood size (1)* | -0.82 | 0.95 | 59 | -0.87 | 0.390 |
|  | Brood size (2) | 0.53 | 0.18 | 60 | 2.94 | 0.005 |
|  | Female age | 0.68 | 0.11 | 60 | 6.08 | <0.001 |

\* Factor removed in the model simplification process. Estimates derived from the last model the factor was included in.

## Discussion

The main aim of this study was to explore whether offspring begging has an influence over parental life-history trade-offs. We found that the experimentally manipulated levels of larval demand influenced the trade-off between number and size of offspring produced (Table 1), but did not directly incur a reproductive cost to the female parent (Table 2, Figure 2a-c). However, we found that the success of the first reproductive event affected the future fecundity of the female negatively (Figure 2e, Table 2). This is the first time that the influence of offspring behaviour on life-history trade-offs has been studied in a similar manner to parental traits (Bryant, 1979; Harding et al., 2009; Hegemann et al., 2013; Tieleman et al., 2008) or manipulations of offspring number (Hodges et al., 2015; Kölliker et al., 2015). Below, we will discuss our findings in more detail.

Our results show that in *N. vespilloides*, the success of the first brood incurs a reproductive cost in fecundity, but not in the number of larvae dispersing from future broods, or in the survival of the parent. Similar effects on fecundity induced by manipulations of parental effort have been implied in previous work: Parejo and Danchin (2006) found that blue tits (*Parus caeruleus*) were less likely to undertake a second reproduction when they had been rearing enlarged broods, but found no evidence for other reproductive costs. Golet et al. (2004) found that black legged kittiwakes (*Rissa tridactyla*) were more likely to breed next year if their eggs were removed the year before, finding that the fecundity costs, when they appeared, were cumulative and appeared in multiple traits. Kölliker et al. (2015) found that the genetic trade-offs between current and future reproduction were more severe before hatching than after hatching. All of these studies imply a difference between the expression of reproductive costs in pre-hatching traits, such as fecundity, and other traits, with the other traits sometimes following a similar pattern to the initial costs apparent at the pre-hatching stage (Golet et al., 2004; Kölliker et al., 2015), and sometimes showing no other costs at all (Parejo and Danchin, 2006). Reproductive costs on fecundity in terms of the number of zygotes produced have been studied less than the number of recruiting offspring, partly due to the majority of data being generated on cavity nesting birds, for which counting eggs imposes bigger methodological difficulties than monitoring recruits. Therefore, data on fecundity costs in the number of zygotes produced, is still lacking, and their existence may well be more common than currently reported data suggests. These initial production costs may very well show the reproductive costs more readily, as they are not under the influence of parental behaviours or conflated by the fitness of the second generation.

In our data, we found a strong correlation between the number of eggs laid and the number of larvae dispersing from the second broods, but only the number of eggs laid was affected by the success of the first broods. The disparity between the two results may be due to parental care masking the costs at the post-hatching stage of juvenile development. In *N. vespilloides*, a similar effect has previously been reported on the effects of egg size on larval size at dispersal: Monteith et al. (2012) found that the size of the larvae at dispersal was determined by the size of the eggs in broods that were raised without parents, but not in ones with them. Thus, it is possible that parental care can also buffer against the post-hatching reproductive costs. Alternatively, the lack of an apparent reproductive cost in the number of larvae at dispersal, can also be tied to the self-feeding ability of *N. vespilloides* larvae (Eggert et al., 1998). The ultimate success of the larvae is a combination of both parental care and their own ability, and the two may also be linked: Providing care for the burying beetle larvae may also improve the larval ability to self-feed.

We found differences in the number and size of offspring produced in the high demand treatment, but not in the other treatments. The difference between this treatment and the intermediate and low demand treatments, is that the level of parental effort is expected to increase rather than reduce in comparison to the natural setting (control treatment). Thus, our result resembles that of a meta-analysis on birds, which reported that reproductive costs on survival tended to only be expressed when the levels of parental effort were increased rather than reduced (Santos and Nakagawa, 2012). The hypothesised positive effects of an energy surplus in treatments where parental effort is reduced from the normal setting (Intermediate and Low treatments), are potentially less plausible, as the females are already likely to be allocating an optimal amount of resources into their current reproduction. Therefore the resources saved from current reproduction may well be invested in other traits, such as those improving the likelihood of gaining another mating opportunity, rather than increasing the number or size of offspring produced during the second reproduction, thus eradicating any benefits that the decreased effort in the first reproductive event could have provided. Male birds have been found to increase their attractiveness when parental effort in previous reproductive event was reduced (Gustafsson et al., 1995; Siefferman and Hill, 2005). However, conclusive determination of where the excess resources are used in *N. vespilloides* requires further experimental work.

We found a positive relationship between the number of larvae produced in the second broods and the lifespan of the females (Table 2). Previous studies investigating the life-history trade-offs have often reported positive relationships between parental effort and life-history traits, rather than the expected negative trade-offs (Reznick, 1985; Roff and Fairbairn, 2007). These positive relationships have often been considered to be a result of differences in individual quality within the study population, as the good quality individuals invest more into a variety of life-history traits connected to fitness and low quality individuals invest less to the same traits (Roff and Fairbairn, 2007). These differences in individual quality can thus mask the underlying trade-off, if there is one (see Wilson and Nussey, 2010). The lack of a trade-off may also be due to differences in the expression of survival effects based on the sex of the parent (Santos and Nakagawa, 2012). Therefore, as we only used females in this study, it is possible that while the burying beetle females did not suffer from survival costs, the males potentially might have. However, since the relationship between the success of the second broods and survival was positive in stead of neutral, the likeliest explanation for the relationship is differences in individual quality of the females.

Lastly, we also found that the overall shape of relationship between the number and size of offspring was non-linear, with an increase at the brood sizes below 20 larvae, after which the average mass of an individual larva in a brood steadily decreases as the brood size increases (Figure 1a). Previous work on *N. vespilloides* and other species of the same genus have reported the classic negative, linear trade-off curves between brood size and the size of the larvae, with the mass of the offspring decreasing with increasing brood size (Smiseth et al., 2014; Trumbo, 1990). However, clutch size and egg volume positively correlated in the same species (Mäenpää and Smiseth, 2017). Resource availability has been shown to have an influence on the shape of the trade-off between number and size of offspring in *Nicrophorus* species before, as the linear relationship has either been shown to be steeper on smaller carcasses (Bartlett and Ashworth, 1988; Scott and Traniello, 1990; Smiseth et al., 2014), or to only affect the number of larvae produced, rather than their size as predicted by the classic life-history theory (Bartlett and Ashworth, 1988; Smith and Fretwell, 1974; Trumbo, 1990; Wilson and Fudge, 1984). The trade-offs have also been shown to only be apparent at the dispersing larvae stage, rather than at the egg production stage, meaning that parental care seems to drive the existence of this trade-off (Mäenpää and Smiseth, 2017; Monteith et al., 2012). Virgin beetles tend to produce approximately 0.8 - 1.4 offspring per a gram of resource on their first reproductive event (reported for *Nicrophorus tomentosus*; Trumbo, 1990; and a similar relationship observed for *N. vespilloides*; personal observation M.I. Mäenpää). Because of this, it can be assumed that the broods are only limited by resources after a certain number of offspring have been produced, the treshold for which is dependent on the size of the carcass. Therefore, we can assume that the broods that are smaller than the optimal size based on the size of carcass available, are not limited by resources, and thus there can be an increase in both the number and size of offspring produced up to the point when the carcass size becomes limiting. After that, the trade-off between number and size can operate.

In conclusion, we found that changes in the level of offspring demand had no effect on the life-history trade-offs of the parent. While there was an overall fecundity cost for the general success of the first broods, the females were able to buffer against this effect at later stages through parental care, without influencing their future survival. As parents that produced more offspring also survived better, the individual quality of the parent plays a role in the resolution of these life-history trade-offs. Individual quality of the female is thus presumably a more important trait in resolving resource allocation between and within broods than offspring demand. However, as offspring demand does have some impact on the success of the current brood, it is nevertheless important to consider its role in changing the nuances behind life-history trade-offs when they do occur. All in all, larval begging does not incur costs to the parents in long term, suggesting that the parents are capable of responding to beggin plastically, and the plastic response has very little influence on the overall life-history of the parent.

